# Single-cell and spatial transcriptomics of the avian embryo tailbud

**DOI:** 10.1101/2024.02.05.578917

**Authors:** GF Mok, S Turner, E Smith, L Mincarelli, A Lister, J Lipscombe, V Uzun, W Haerty, IC Macaulay, A Münsterberg

**Author notes:** Department of Zoology, University of Cambridge, Downing Street, Cambridge CB2 3EJ, UK. Wellcome Sanger Institute, Wellcome Trust Genome Campus, Hinxton, Saffron Walden CB10 1RQ, UK.

## Abstract

Vertebrate body axis formation initiates during gastrulation and continues within the tail bud at the posterior end of the embryo. Major structures in the trunk are paired somites, which generate the musculoskeletal system, the spinal cord - forming part of the central nervous system, and the notochord, with important patterning functions. The specification of these different cell lineages by key signalling pathways and transcription factors is essential, however, a global map of cell types and expressed genes in the avian trunk is missing. Here we use single-cell RNA sequencing and RNA tomography to generate a molecular map of the emerging trunk and tailbud in the chick embryo. Single cell RNA-sequencing (scRNA-seq) identifies discrete cell lineages including somites, neural tube, neural crest, lateral plate mesoderm, ectoderm, endothelial and blood progenitors. In addition, high-throughput RNA-seq of sequential tissue sections provides a spatially resolved, genome-wide expression dataset for the avian tailbud and emerging body, comparable to other model systems. Combining the single-cell and spatial datasets, we identify spatially restricted genes, focusing on somites and early myoblasts. Thus, this high-resolution transcriptome map incorporating cell types in the embryonic trunk can expose molecular pathways involved in body axis development.

## Introduction

The generation of somites, which arise in a regular sequence during embryogenesis, is fundamental for creating the vertebrate segmented body plan (Benazeraf and Pourquie, 2013). Pairs of somites form on either side of the neural tube from unsegmented, paraxial mesoderm, and the process of somitogenesis, which involves waves of cycling gene expression, has been studied extensively in chick embryos (Pourquie, 2004). Prospective paraxial mesoderm cells emerge from the primitive streak during gastrulation (Psychoyos and Stern, 1996) and follow a stereotypical migration trajectory towards their destination (Iimura et al., 2007; Yang et al., 2002). As the body axis elongates, bi-potential neuro-mesodermal progenitors (NMP) located in the tailbud continue to generate paraxial mesoderm and cells of the neural tube (Henrique et al., 2015; Wilson et al., 2009; Wymeersch et al., 2021). The dynamics of this specialised cell population has been mapped in detail in chick embryos (Guillot et al., 2021) and it has been shown that the extension of neural and paraxial mesoderm tissues in the embryonic body is coordinated by mechanical interactions (Xiong et al., 2020).

Somite differentiation proceeds along the posterior-to-anterior axis and serves as a paradigm for the study of cell fate specification. Multiple signals from surrounding tissues are integrated by somite cells to produce the lineages of the musculoskeletal system, including chondrocytes of the axial skeleton and skeletal muscles of the trunk and limbs (Brent and Tabin, 2002; Christ et al., 2007; Christ and Scaal, 2008). Cell fate specification is intimately linked to stereotypic morphological changes resulting in somite compartmentalisation. Tracking of GFP-labelled cells showed that the dorsal dermomyotome produces the myotome layer in multiple waves, with the first myocytes specified adjacent to the neural tube (Gros et al., 2004). Live-imaging of cellular rearrangements examined the morphological transformations of somites from epithelial structures to somites with a mesenchymal sclerotome, located ventrally, and an epaxial myotome abutting the neural tube (McColl et al., 2018). This uncovered differential cell sizes and regions of proliferation as well as a directed movement of dermomyotomal progenitor cells towards the rostro-medial domain of the dermomyotome, where skeletal muscle formation initiates.

To better characterise the regulation of these morphogenetic events and their integration with cell specification and differentiation, we previously assessed the dynamic changes of the transcriptome and of chromatin accessibility across presegmented mesoderm and early somites (Mok et al., 2021). Associating differentially accessible chromatin with nearby genes differentially expressed along the axis identified candidate cis-regulatory elements (CREs) involved in expression of transcription factors important for somite formation and differentiation. Time-lapse microscopy in accessible chick embryos of fluorescent CRE-reporters revealed their spatio-temporal activity and mutation analysis uncovered some upstream regulators. Similarly in mice, we examined matched gene expression and open chromatin profiles for newly formed somite pairs across a developmental time series. This provided a high-resolution view of the molecular signatures underlying the conserved maturation programme followed by all somites after segmentation (Ibarra-Soria et al., 2023).

Here we focus on the initial phase of trunk development in chick embryos. Recent in vitro organoid models of axis elongation based on the differentiation of mouse or human pluripotent stem cells use pharmacological activation or inhibition of crucial signalling pathways (Veenvliet and Herrmann, 2021). These self-organising structures include gastruloids (Moris et al., 2020; Turner et al., 2017), trunk-like structures (Veenvliet et al., 2020), or somitoids (Sanaki-Matsumiya et al., 2022), or axioloids (Yamanaka et al., 2023). These often comprise mesoderm, including somites, although the notochord, which is involved in patterning of trunk tissues, is missing. Thus, it is important to reconstruct the molecular profiles and cellular composition in the native tissues.

Here, we use single-cell transcriptomics combined with an RNA-tomography based approach, analogous to Tomo-seq (Junker et al., 2014; Kruse et al., 2016) but using a modified G&T-seq approach (Macaulay et al. 2015). This generated a spatio-temporal map of the emerging trunk and identified genes not previously known to be involved in presomitic mesoderm (psm) and somite maturation. Our study complements data in chick embryos, from earlier developmental stages (HH4-HH11) (Rito et al., 2023; Vermillion et al., 2018; Williams et al., 2022), from tailbud (Guillot et al., 2021) and from prospective neural plate, neural plate border and non-neural ectoderm (Trevers et al., 2023). The dataset is relevant for cell-type specification during early body formation and may provide insights into the molecular genetics that underlie diseases of the musculoskeletal system.

## Results

### Single cell profiling of the developing chick embryonic body

To investigate the molecular basis underpinning somitogenesis and axis elongation of the growing chick body, we mapped the transcriptomes of individual cells at embryonic stage HH14 (Hamburger and Hamilton, 1992). This stage embryo has 22 somites, including cervical level somites (6-19) and thoracic level somites (20-22). The unsegmented paraxial mesoderm and tailbud comprise prospective somites of the thoracic, lumbar and sacral regions (Weldon and Munsterberg, 2022). Single cell suspensions from the posterior part of five pooled embryos included the extraembryonic region, tailbud, pre-somitic mesoderm and the most recently formed six somites. Following enzymatic digestion and mechanical dissociation the suspension was processed using the 10X Genomics Chromium (Fig. 1). A total of 6158 cells were sequenced with a median of 517 genes and 900 UMIs per cell.

**Figure 1:**
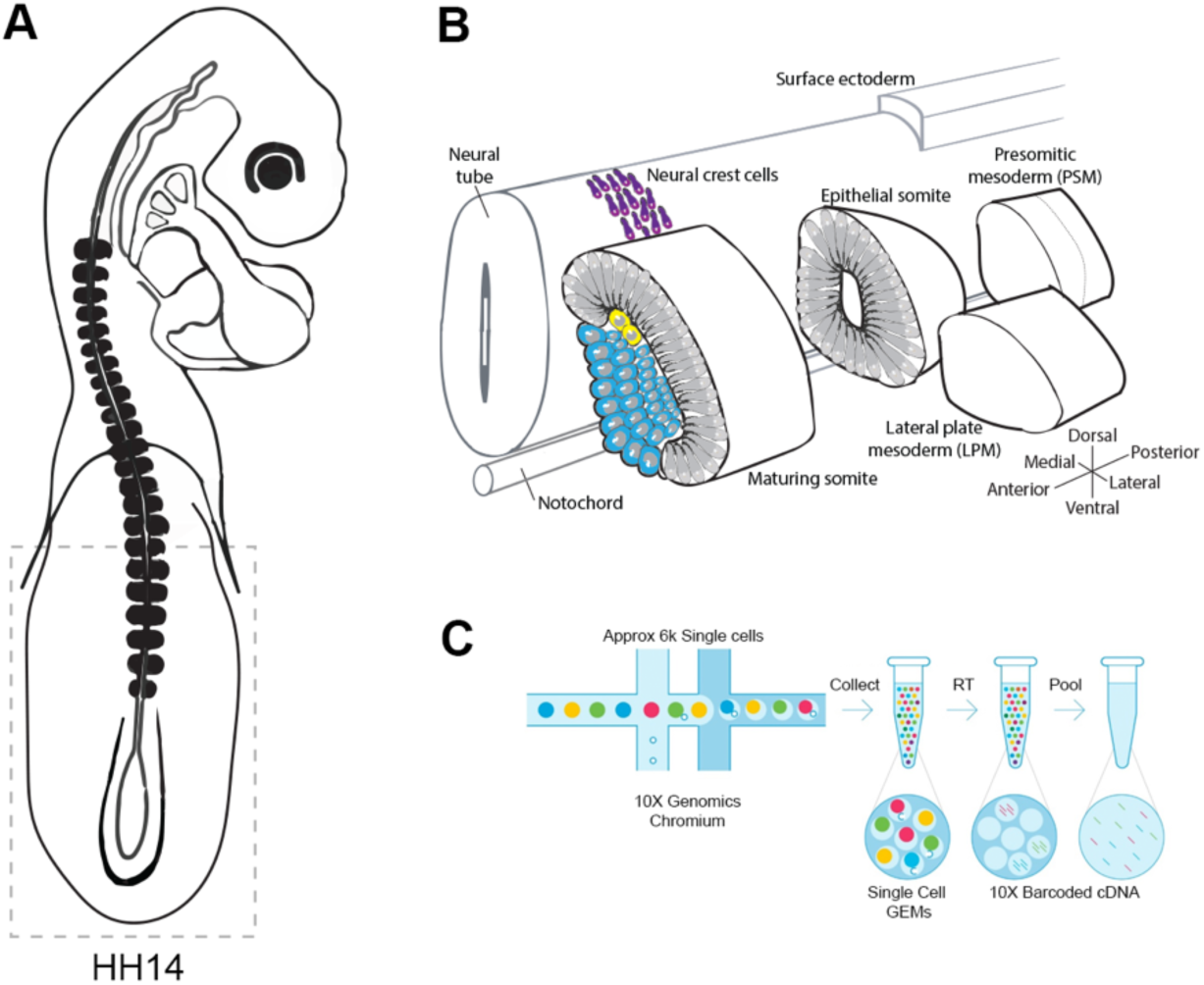
High-throughput scRNA-seq of the embryonic trunk. (A) Trunk regions, indicated by stippled lines, of five stage HH14 chicken embryos were collected for scRNA-seq. (B) Illustration of the developing structures captured: pre-somitic mesoderm (psm), epithelial somites, maturing somites, lateral plate mesoderm (lpm), surface ectoderm, neural crest cells and notochord. (C) Schematic of the 10X Genomics Chromium workflow.

Unsupervised clustering was used to classify cell populations (Butler et al., 2018). Projection onto UMAP plots revealed 10 separate clusters (Fig. 2). Classic marker genes assigned cluster identity and tissues: lateral plate mesoderm (*Prrx1* and *Krt18*), neural (*Wnt4* and *Olig*2), epithelial somites (*Meox1* and *Tcf15*), maturing somite (*Nkx3-2* and *Twist1*), ectoderm (*Fabp3* and *Wnt6*), blood (*Hbm* and *Hba1*), tailbud (*Cdx4* and *Msgn1*), endothelial (*Lmo2* and *Sox18*), notochord (*Tf* and *Shh*) and neural crest cells (*FoxD3* and *Sox10*) (Fig. 2A-C). Seurat cell cycle scoring determined cell cycle activity. This showed cell clustering was not due to cell cycle phase, although neural cells were predominantly in S phase and G2M phase (Fig. 2D).

**Figure 2:**
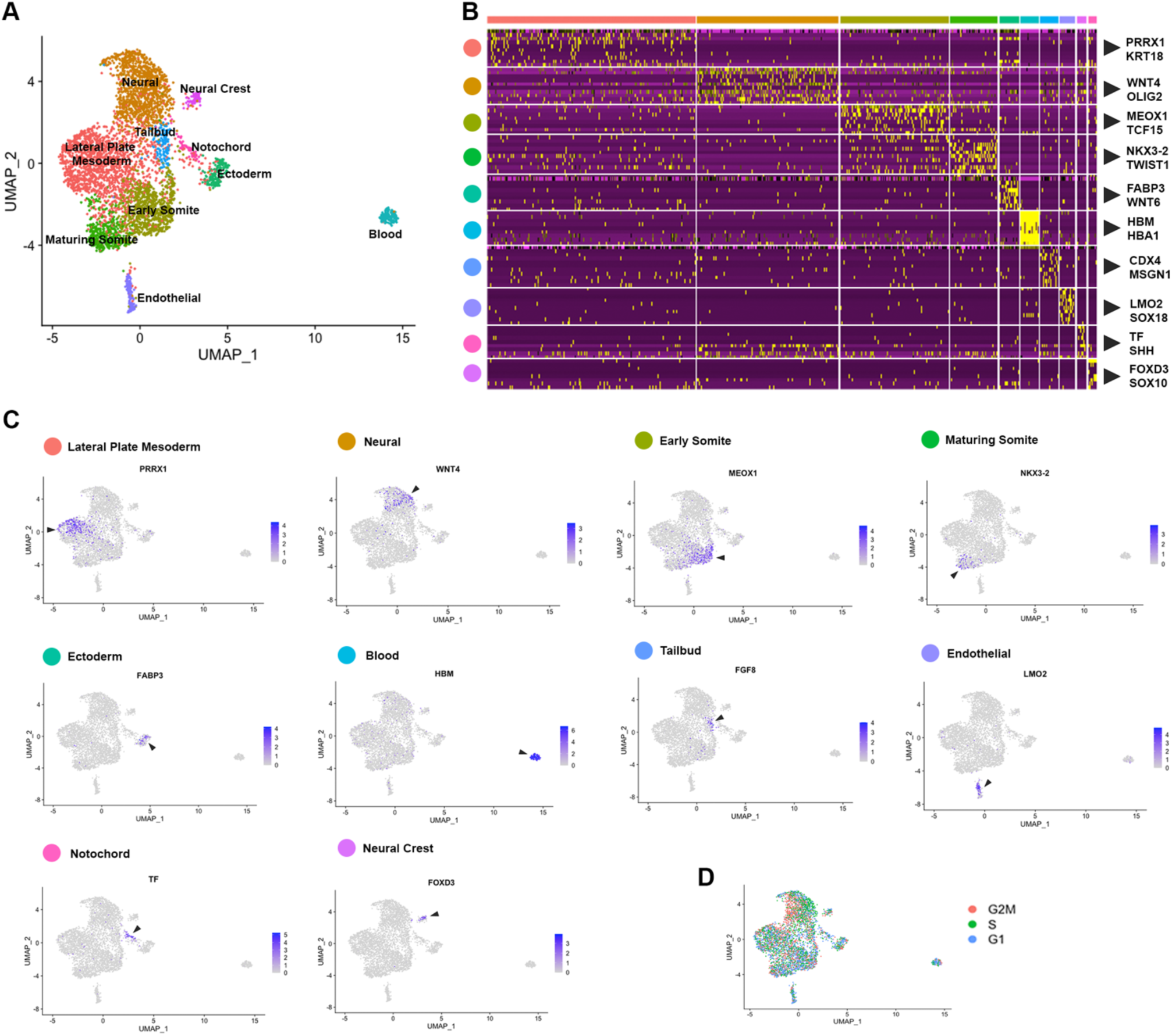
Cell population composition and signatures of the HH14 chicken embryo trunk. (A) Unsupervised UMAP subdivides cells within the trunk into 10 clusters – lateral plate mesoderm, neural progenitors, early somite, maturing somite, pre-somitic mesoderm, ectoderm, blood progenitors, endothelial progenitors and neural crest. (B) Heatmap of the top 10 genes significantly enriched in each cluster; representative genes are shown. (C) UMAPs show log normalised counts of representative genes for each cluster. Colour intensity is proportional to expression level of each gene. (D) Distribution of cell cycle phases visualised using Seurat cell cycle scoring.

### Spatial transcriptomics profiling of the developing chick embryonic trunk

Spatial information is lost in single cell sequencing data following tissue dissociation (Griffiths et al., 2018). To address this, we next used a spatial transcriptomics approach to quantify the transcriptomes of a series of individual cryogenic sections along the HH14 embryonic trunk. This enabled a systematic investigation of spatial RNA profiles along the axis. Libraries were generated from 20 micron consecutive cryosections of a HH14 chick embryo. Each section was collected into lysis buffer and mRNA captured and amplified using a modification of the G&T-seq protocol (Macaulay et al., 2016). The resulting cDNA libraries had high complexity and enabled us to confidently determine spatial gene expression (Fig. 3).

**Figure 3:**
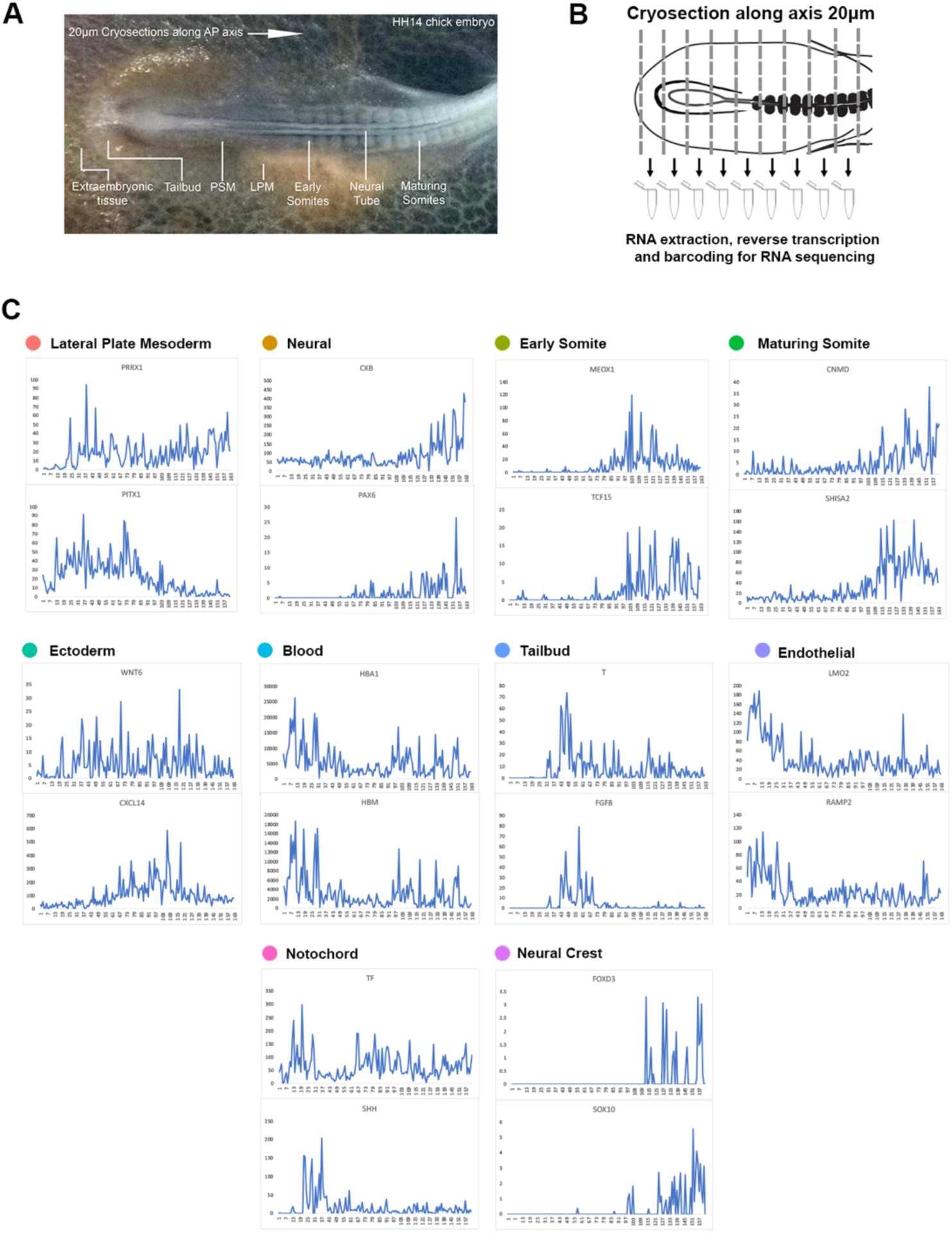
Spatial sequencing reveals distinct gene expression profiles along the embryonic axis. (A) Stage HH14 chick embryo trunk was sectioned along the anterior-to-posterior axis, from extraembryonic tissue at the posterior end, through the tailbud and pre-somitic mesoderm towards maturing somites. Individual sections were collected in wells followed by RNA isolation and cDNA preparation using section specific barcodes. After that, samples were pooled for linear amplification and sequence library preparation. (B) Spatial expression traces for representative genes in each corresponding tissue type identified from the scRNA-seq clustering.

Using the same markers as in the previous scRNA-seq analysis, we identified 10 different clusters and established profiles of the lateral plate mesoderm, neural tissue, early somite, maturing somite, ectoderm, blood, endothelial, tailbud, notochord and neural crest cells (Fig. 3). Spatial patterns of localised anterior-to-posterior restricted gene expression were evident for all tissue types, with exception of the ectoderm, suggesting that this tissue has few distinguishing markers along the A-P axis. In the most posterior samples, we identified blood *(Hbm*, *Hba1*) and endothelial marker genes (*Lmo2*, *Ramp2*), consistent with this region comprising extra-embryonic tissue. Subsequent sections showed the onset of tailbud genes (*T* (= *brachyury*) and *Fgf8*). Some notochord markers were more highly expressed in posterior sections (*Shh*), while transferrin (*Tf*) was expressed along the axis, with lower levels around the tailbud region. Across neural tissue (*Ckb*, *Pax6*), early somites (*Meox1*, *Tcf15*) and maturing somites (*Cnmd*, *Shisa2*), gene expression profiles showed a gradual increase towards the anterior. Neural crest cells are beginning to migrate and become distinguishable in the more anterior sections with discreet profiles detected for *Foxd3* and *Sox10* (Fig. 3C).

### Spatial transcriptomics resolves mRNA localisation patterns

To identify gene expression patterns systematically, we clustered our spatial gene expression data based on a self-organising heatmap. This sorted the cumulative gene expression traces along a linear axis of 180 profiles and identified 3 major groups of localised mRNA (Fig. 4A). The first group localised to the most posterior, the second group displayed an increase in the tailbud region and across the pre-somitic mesoderm, and the third group of transcripts was most highly expressed in the anterior sections comprising epithelial and maturing somites (Fig. 4B). Transcripts enriched posteriorly (e.g. *Hbm*, *Epas1*, *Lmo2*) were related to hematopoiesis, erythrocyte differentiation and myeloid homeostasis, consistent with the presence of extraembryonic blood islands. The second profile showed genes enriched for Gene Ontology (GO) processes such as anterior-posterior pattern specification, embryo morphogenesis and tissue morphogenesis. This overlaps spatially with tailbud and pre-somitic regions. Markers with enriched expression included *Wnt5a*, *Msgn1*, *Tbx6* and *Fgf8*. The third profile, which overlaps with the formation of somites but also neural tube development, included genes enriched for pattern specification, neurogenesis and animal organ morphogenesis, such as *Meox1* and *Shisa2*. Interestingly, the profiles for somites and neural tube are very similar and genes were clustered together, suggesting these tissues mature at a similar rate. However, GO analysis did separate genes associated with either somitogenesis or neurogenesis (Fig. 4C). Spatial transcriptomics also detected the opposing gradients across the psm and somites of *Wnt5a*, *FGF8* and *Aldh1a2*, encoding an enzyme involved in retinoid acid synthesis. The transcripts for *Meox1* and *Tcf15* become upregulated in anterior psm and epithelial somites, whereas *Mesp1* transcripts are restricted to an anterior region in the psm, comprising the next but one prospective somite. Transcripts for *Hoxd11*, *Hoxc9* and *Hoxb1* show the expected expression boundaries along the anterior-posterior axis (Fig. 4D). Analysis of expression levels of components of important signalling pathways, including WNT, FGF, NOTCH and BMP pathways, shows high levels of *Wnt5a* and *Fgf8* in tailbud population, high levels of *Wnt6* in the ectoderm, *Notch1/2* in neural and neural crest cells and *Bmp4* in the lateral plate (Fig. 4E).

**Figure 4.**
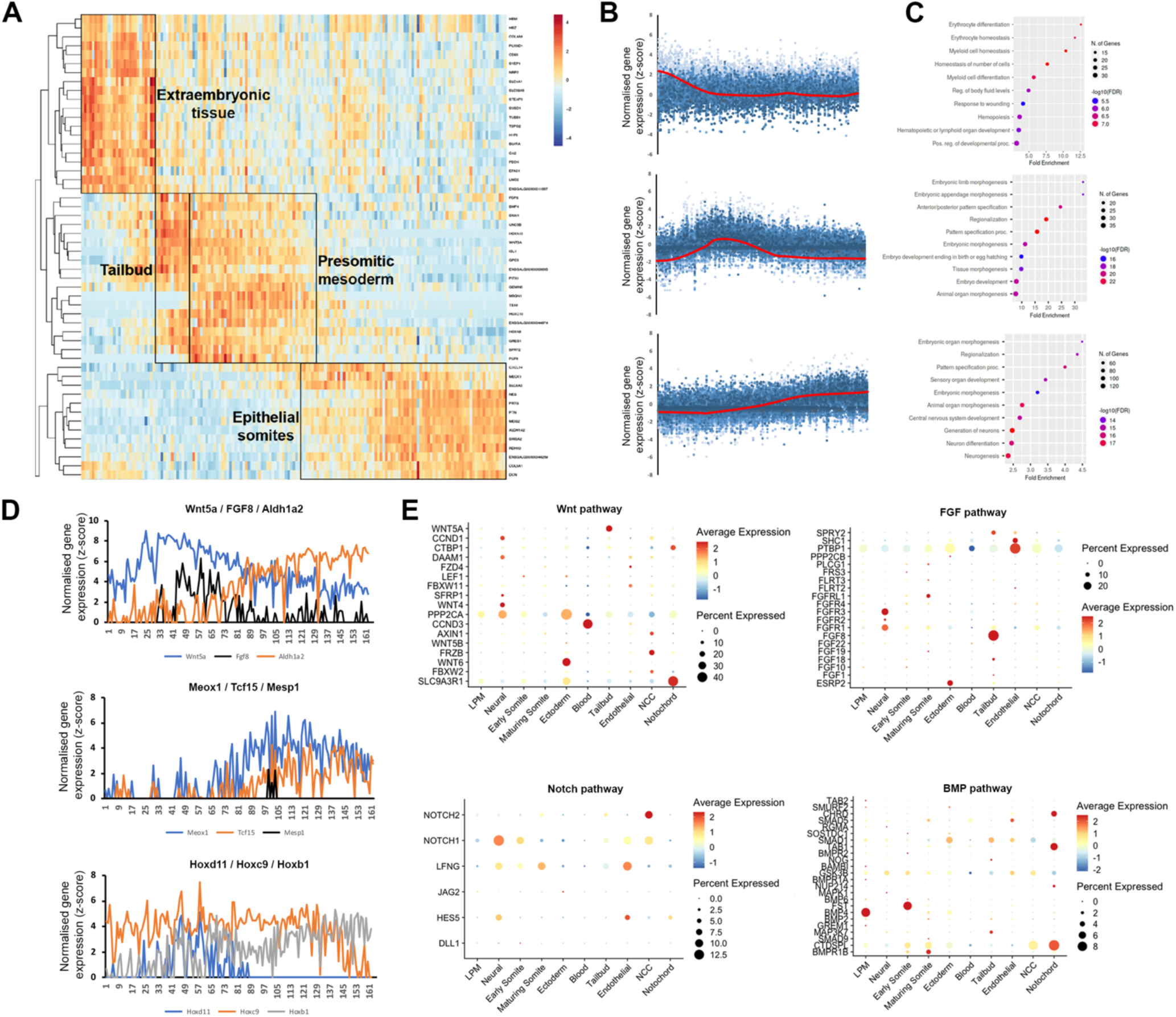
K-means clustering identifies biological components along the posterior-to-anterior axis. (A) Hierarchical cluster analysis of gene expression per section (total 180). Distinct gene expression clusters correspond to different regions along the axis, characterised by extraembryonic tissue, tailbud, pre-somitic mesoderm and epithelial somites – indicated by boxed areas. RNA sequencing reads per gene were normalised against the total read count per section. (B) Spatial expression traces for representative genes in each corresponding cluster. (C) Gene ontology on genes enriched in the extraembryonic tissue, tailbud and pre-somitic mesoderm, and epithelial somites and neural tube. (D) Spatial expression traces for signalling pathways associated with anterior-posterior patterning such as WNT (*Wnt5A*), FGF (*Fgf8*) and retinoic acid (*Aldh1A2*); for genes associated with paraxial mesoderm differentiation (*Meox1*, *Tcf15* and *Mesp1*); and for *Hox* genes involved in anterior-posterior patterning (*Hoxd11*, *Hoxc9* and *Hoxb1*). (E) Dot plot showing average expression of genes and percentage expressed in each cell cluster associated with the WNT, FGF, NOTCH and BMP signalling pathways.

### Correlation of single cell RNA sequencing with spatial transcriptomics

Axis patterning is characterized by the progressive differentiation of cell types with anterior-to-posterior identity. To validate genes identified in specific clusters obtained from the single-cell RNA-sequencing, we correlated spatial patterns and confirmed gene expression by *in situ* hybridisation. Expression of *Hbm*, a haemoglobin gene, is representative for the most posterior group of transcripts, which is enriched for genes involved in hematopoietic differentiation. We show that *Hbm* transcripts are restricted to blood islands in posterior extraembryonic tissue (Fig. 5A). The second spatial cluster values correlate with the tailbud and pre-somitic mesoderm. We validated this data using *Wnt5a*, a known regulator of cell movement behaviour during axis elongation (Sweetman et al., 2008). *In situ* hybridisation showed *Wnt5a* expression was restricted to the tailbud region (Fig. 5B), where neuromesodermal progenitor (NMP) cells are located. The third spatial cluster, where gene expression increased towards the anterior, was validated using *Tbx22* - a key gene for somite boundary formation. Expression of *Tbx22* was restricted to caudal somite domains. The periodicity visualised *in situ* correlates with the distribution in the UMAP and in the spatial line plots (Fig. 5C).

**Figure 5:**
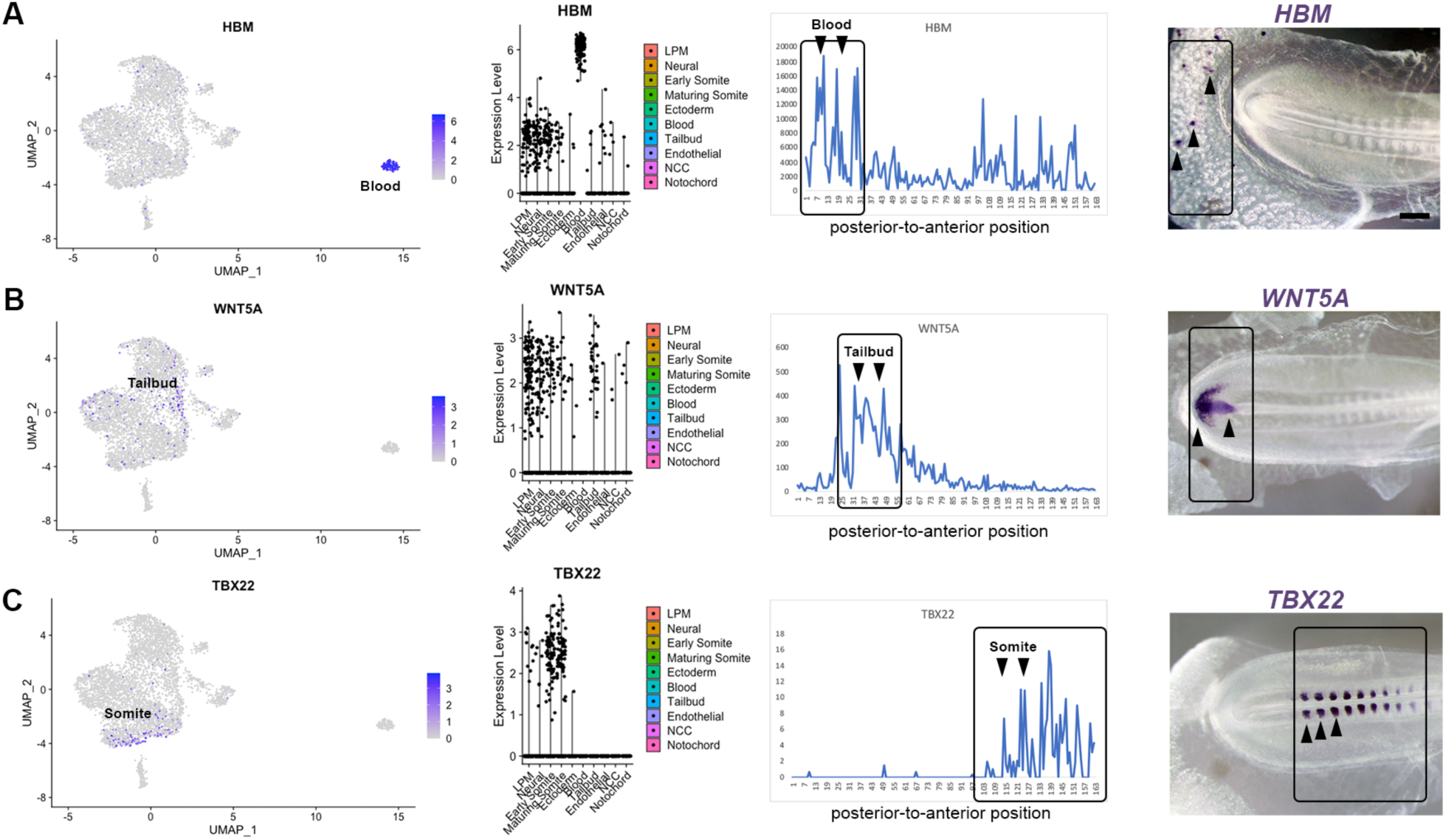
Transcriptome map of the embryonic trunk with high spatial resolution. (A-C) UMAP plot and violin plot of representative genes for different tissues, with comparison to Spatial expression trace along the posterior-to-anterior axis. The corresponding in situ hybridisation is shown for (A) *Hbm* – blood islands, (B) *Wnt5A* - tailbud, (C) *Tbx22* – caudal somite halves.

### Identification of new genes in paraxial mesoderm and somites

Next, we investigated previously unexplored genes uncovered by our approach. We focused on candidates with restricted expression in the pre-somitic mesoderm and somites. Using the subset feature in Seurat, we profiled three clusters from the single-cell RNA seq dataset – tailbud, early somite and maturing somite (Fig. 6A). We re-ran the clustering, findneighbours and pca tests on this new Seurat object. Classic markers for each cluster were re-plotted on UMAPs, such as *Msgn1* for the pre-somitic mesoderm, *Meox1* and *Tcf15* for early somites and *Tbx22* for maturing somites (Fig. 6A). Sub-clustering also revealed restricted expression of follistatin (*Fst*), in a group of cells potentially representing early myoblasts located adjacent to the neural tube. We identified three genes for further analysis. UMAP plot of *Olfml3* and *Foxd1* gene expression suggested they are likely to be expressed in maturing somites, whereas *Lrig3* was predicted to be expressed in some pre-somitic mesoderm cells, early somites and less in maturing somites (Fig. 6A).

**Figure 6:**
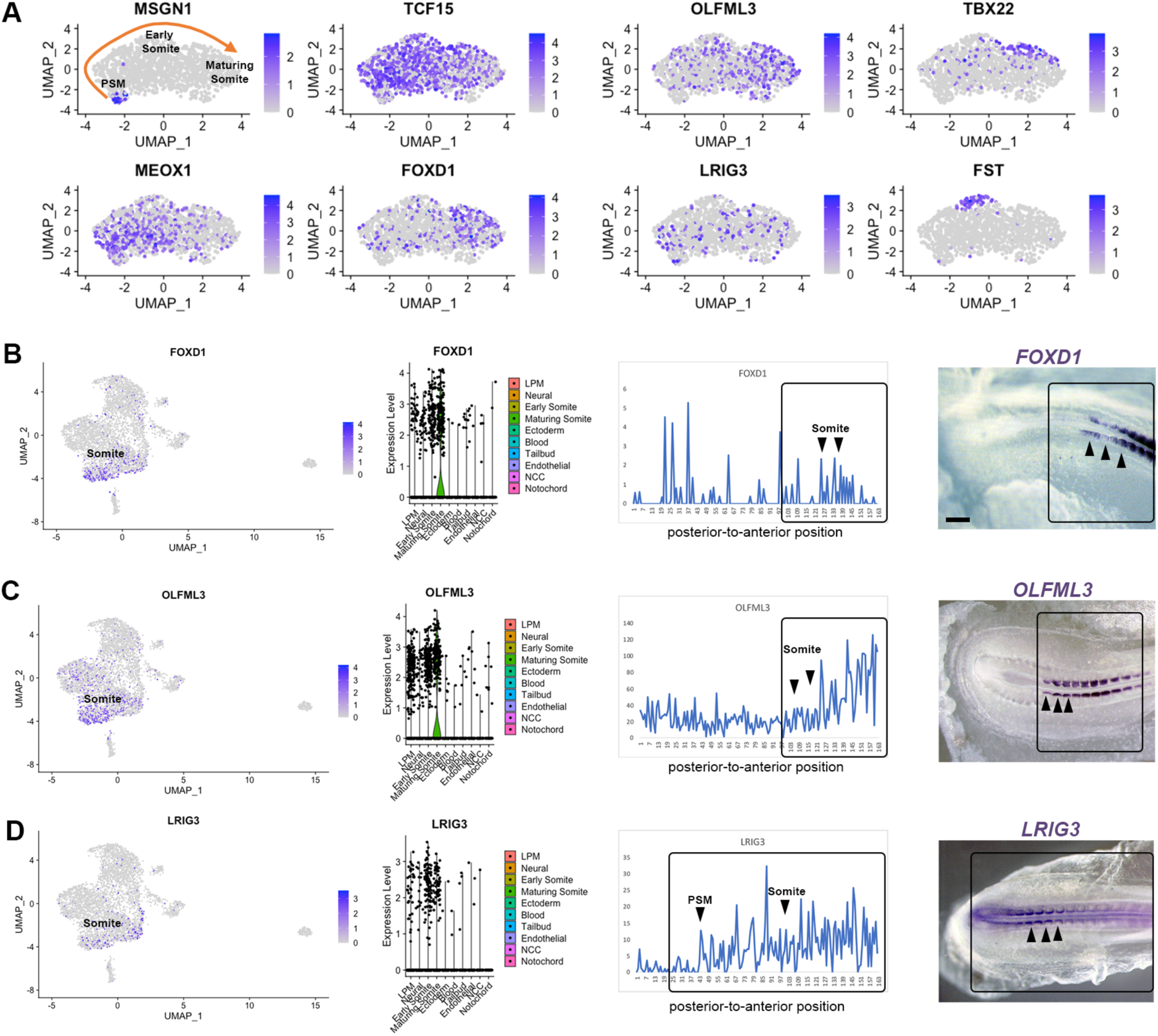
Differentially expressed genes in pre-somitic mesoderm and somites. (A) Sub-clustering identifies specific genes within the psm, early somites and maturing somites. Psm is characterised by *MSGN1* expression while *Tcf15*, *Meox1* and *Tbx22* markers represent somites. Restricted *Fst* expression may correlate with epaxial myoblasts. (B-D) UMAP, violin plots and spatial expression traces show expression for *Foxd1*, *Olfml3* and *Lrig3*, not previously identified in somites. Whole-mount in situ hybridisation confirmed the spatially restricted expression of *Foxd1* and *Olfml3* in epithelial and maturing somites and of *Lrig3* in the psm and somites.

Interrogation of the profiles for these genes in the spatial data showed that expression of *Foxd1* and *Olfml3* increased towards the anterior regions. *Foxd1* and *Olfml3* were also identified in spatial cluster 3 (Fig. 4). For *Lrig3*, the spatial data showed increasing expression from the most posterior to the anterior embryonic regions, suggesting that it is expressed from the tailbud to maturing somites. Spatial validation for all three genes *in situ* confirmed these observations: *Foxd1* and *Olfml3* are restricted to somites whilst *Lrig3* is expressed in the tailbud, pre-somitic mesoderm and in somites. In epithelial somites, all three genes are restricted medially. In maturing somites *Foxd1* is broadly expressed, *Olfml3* remains medially restricted and *Lrig3* is downregulated (Fig. 6B-D).

## Discussion

The chick embryo is a classic model for developmental biology studies due to the versatility of *in vivo* experimental approaches (Gandhi and Bronner, 2018; Sauka-Spengler and Barembaum, 2008; Stern, 2005). For example, it has served to better understand the processes of body axis formation, segmentation/somitogenesis and differentiation (Benazeraf and Pourquie, 2013). Here we use single cell transcriptomics and spatial transcriptomics to map cells that arise in the emerging trunk as it extends. This adds to the growing body of literature, which includes scRNA-seq data of the chick embryo from primitive streak to neurula stages (Vermillion et al., 2018; Williams et al., 2022). Previous work characterised the molecular signature of neuromesodermal progenitors (NMP) in detail by micro-dissecting anterior PS in stage HH5 and in 6-somite embryos (HH9^-^) as well as the tail bud of 35-somite embryos (∼HH18) (Guillot et al., 2021). A recent report examined the anterior most part of the main body axis, including occipital and cervical somites at several stages of development, 4-somites, 7-somites, 10-somites and 13-somites (Rito et al., 2023). This work is in agreement with our data presented here. We identified similar cell populations when the body extends and the cervical-thoracic region forms (HH14, 22-somite embryo) (Weldon and Munsterberg, 2022).

Furthermore, we show that combining scRNA-seq with spatial transcriptomics can reveal novel genes involved in specific aspects of axis extension. As an example, we focused on paraxial mesoderm and discovered the previously unknown relevance of *Foxd1*, *Olfml3* and *Lrig3* in developing somites. All three genes showed restricted expression in the medial somite domain suggesting a possible role in early myoblasts. It is noteworthy that *Foxd1*, a member of the fork-head family of transcription factors, is associated with pluripotency and seems to be required for successful reprogramming (Koga et al., 2014). In addition, *Foxd1* protects senescence in human mesenchymal stem cells (hMSC) and is regulated by YAP (Fu et al., 2019). Not much is known about *Olfml3* function in development. It is a secreted glycoprotein of the Olfactomedin-family, which organises the extracellular matrix and has pro-angiogenic properties. *Olfml3* deficient mice exhibit abnormalities in the vasculature causing lethality (Imhof et al., 2020). *Olfml3* has also been implicated in pre-natal muscle development in pig (Jin and Li, 2019) and in *Xenopus* it is involved in dorso-ventral patterning by enhancing chordin degradation (Inomata et al., 2008). Finally, Leucine-rich repeats and immunoglobulin-like domains 3 (*Lrig3*) plays a role in neural crest development (Zhao et al., 2008) in *Xenopus*. This is consistent with studies in mice, which showed *Lrig3* is involved in inner ear morphogenesis by restricting the expression of *Ntn1* (Abraira et al., 2008). However, the roles of these genes in developing somites have not yet been investigated.

Tomo-seq is a spatial transcriptomics approach first used in zebrafish embryos (Junker et al., 2014; Kruse et al., 2016). We modified and automated this approach using the G&T-seq protocol (Macaulay et al. 20015) and applied it to the posterior half of a HH14 whole embryo. As reported previously in the zebrafish heart (Burkhard and Bakkers, 2018; Wu et al., 2016), we obtained high spatial resolution and sensitivity as shown by hierarchical cluster analysis. Known marker genes were expressed in the anticipated spatio-temporal patterns and identified the appropriate regions along the body axis. The extraembryonic tissue was characterized by *Hbm*, *Hbz*, *Lmo2* and *Epas1*, the tailbud region by *Hoxa13*, *Wnt5A*, the presegemented mesoderm by *Msgn1*, *Tbx6* and epithelial somites by *Meox1* and genes involved in retinoic acid (RA) signalling.

Overall, this study integrates scRNA-seq with spatial transcriptomics improving our understanding and validating the gene expression patterns within the avian tailbud. While the scRNA-seq provides information on gene expression at the single-cell level, combining it with spatial transcriptomics using the G&T method allows preservation of the spatial context of gene activity along the anterior-to-posterior axis. Although other technologies are now available for obtaining high-resolution spatial transcriptional profiles of tissues, such as MERFISH, 10X Genomics Visium and Xenium, these would be costly to implement to understand gene dynamics along the anterior-to-posterior axis with multiple samples. Our results illustrate the advantage of combining different approaches to address fundamental questions in developmental biology.

## Materials and Methods

### Chicken embryos

Fertilised chicken eggs (Henry Stewart & Co.) were incubated at 37 °C with humidity. Embryos were staged according to (Hamburger and Hamilton, 1992). All experiments were performed on chicken embryos younger than two thirds of gestation and therefore were not regulated by the Animal Scientific Procedures Act 1986.

### Preparation of single cells from chicken embryos

The trunk of HH14 embryos were dissected into Ringer’s solution in silicon lined petri dishes and pinned down using the extra-embryonic membranes. Embryonic tissue was transferred into low binding tubes and Ringer’s solution was replaced with Dispase (1.5 mg/ml) in DMEM 10 mM HEPES pH7.5 at 37 °C for 7 min prior to treatment with Trypsin (0.05%) at 37 °C for 7 min. The reaction was stopped with Ringer’s solution with 0.25% BSA. Cells were spun down and resuspended in Hank’s solution prior to passing through 40 μm cell strainer to obtain single cell suspension.

### scRNA-seq library preparation

A suspension of approximately 10,000 single cells was loaded onto the 10X Genomics Single Cell 3’ Chip. cDNA synthesis and library construction were performed according to the manufacturer’s protocol for the Chromium Single Cell 3’ v2 protocol (PN-12033, 10X Genomics). Samples were sequenced on Illumina HiSeq 4000 100 bp paired-end runs.

### scRNA-seq data analysis

Cell Ranger 3.0.2 (10X Genomics) was used to de-multiplex Illumina BCL output, create fastq files and generate single cell feature counts for the library using the *Gallus gallus* transcriptome (Ensembl release 94) with 82.4% reads mapped to the genome. Subsequent processing was performed using the Seurat v3.1.0 (Butler et al., 2018) package within R (v3.6.1). Cell quality was assessed using simple QC metrics: total number of expressed genes, mitochondrial RNA content and ribosomal RNA content. Identification of chicken mitochondrial RNA content and ribosomal RNA content. Outlier cells were identified if they were above or below three median absolute deviations (MADs) from the median for any metric in the dataset. Data was normalised across all cells using the ‘LogNormalize’ function with a scale factor of 1e4. A set of genes highly variable across the cells was identified using the ‘FindVariableGenes’ function (using ‘vst’ and 2000 features) before being centred and scaled using the ‘ScaleData’ function with default parameters. PCA analysis was performed on scaled data using variant genes and significant principal components were identified by plotting the standard deviation of the top 50 components. The first 2 principal components showed high enrichment for mitochondrial genes and were subsequently regressed and only principal components 3:25 were used to create a Shared Nearest Neighbour (SNN) graph using the ‘FindNeighbours’ function with k.param set to 10. This was used to identify clusters of cells showing similar expression profiles using the FindClusters function with a resolution set to 0.6. The Uniform Manifold Approximation and Projection (UMAP) dimensional reduction technique was used to visualise data from principal components 3:26 in two-dimensional space (‘RunUMAP’ function). Graphing of the output enabled visualisation of cell cluster identity and marker gene expression. Biomarkers of each cluster were identified using Wilcoxon rank sum tests using Seurat’s ‘FindAllMarkers’ function. It was stipulated that genes must show a logFC of at least 0.01 to be considered for testing. Only positive markers were reported. The expression profile of top markers ranked by average logFC were visualised as heatmaps and dotplots of the scaled data. Cluster identity was determined using visual inspection focussing on the expression of known marker genes. For cells identified in tailbud and somite clusters, the ‘Subset’ function was used to create a new Seurat object which narrowed down to 1359 cells. Using the ‘FindNeighbors’ feature with dimensions set to 3:20, ‘FindClusters’ resolution of 0.4 and ‘RunUMAP’ set with dimensions 3:16.

### Spatial RNA sequencing from sections

Embryos were embedded in Jung tissue freezing medium (Leica), orientated and rapidly frozen on dry ice, and stored at −80° C prior to cryosectioning. Embedded embryos were cryosectioned at 20 μm thickness, collected into 96-well plates (on ice) prior to the addition of 10 μL of RLT plus lysis buffer (Qiagen, Hilden, Germany). All instruments and surfaces were cleaned with 80% v/v ethanol, RNAse-free water and lastly RNAse-out solution after each sample to reduce cross-contamination and RNA degradation. Samples were stored at −80° C until cDNA preparation using the G&T-seq method as previously described (Macaulay et al. 2015) with minor modifications to accommodate the larger volume of lysis buffer. cDNA was normalised to 0.2 ng/μL before Nextera (Illumina, San Diego, CA, USA) library preparation in a total reaction volume of 4 μL. Libraries were pooled by volume and sequenced on a single lane on the Illumina HiSeq 2500 (150-bp paired-end reads).

### Low-input RNA sequencing analysis

For RNA-seq analysis, we used Refseq version GRCg6a for genome assembly and gene annotation. Reads were trimmed and adapters were removed using tim-galore version 0.4.2. Heatmap based on hierarchical clustering was generated in R-Studio version 1.2.1335 and plotted as a heatmap using the R package DeSeq2 (Love et al. 2014).

### Wholemount in situ hybridisation

Wholemount in situ hybridisation using DIG-UP labelled antisense RNA probes was carried out using standard methods. Probes were generated from amplificons of chicken cDNA using the following primers: *Wnt5a* (GCAGCACTGTGGACAACAAC/CACCGTCTTGAACTGGTCGT), *Olfml3* (GGGAGTTCACGCTCTTCTCG/GATGATCTGGTAGCCGTCGT) *Hbm* (CATCACACATTGCCACCAG C/GCAGCAATGGTGTCTTTATTGA), *Tbx22* (GGATGTTCCCATCGGTCAGG/AGACTTAGCGCTCTT CAGGC), *Lrig3* (GTCCTGACGCCTGGGAATTT/AATCTGTGGGACAGGATGCC). Briefly, following fixation in 4% PFA embryos were treated with Proteinase K, hybridised with the probe over night at 65°C. After post-hybridisation washed and blocking with BMB (Roche), embryos were treated with anti-DIG antibody coupled to alkaline phosphatase (Merck) and signal developed using NBT/BCIP (Melfords Laboratories).

## Data availability

For scRNA-seq the raw sequencing data can be accessed on the NCBI-SRA archive under accession number tbc. Raw spatial transcriptomics data are available under NCBI-SRA accession number tbc.

## ACKNOWLEDGEMENTS

The authors acknowledge support from the Biotechnology and Biological Sciences Research Council (BBSRC), part of UK Research and Innovation, Core Capability Grant BB/CCG1720/1 and the National Capability BBS/E/T/000PR9816 (NC1 - Supporting EI’s ISPs and the UK Community with Genomics and Single Cell Analysis). ICM was supported by a BBSRC New Investigator Grant BB/P022073/1. AM acknowledges funding from BBSRC to support GM (BB/N007034/1) and a NRPDTP studentship to support ES. ST was supported by a Summer internship held at EI.

## AUTHOR CONTRIBUTIONS

Conceptualization: GM, AM, WH and IM. Funding acquisition, Project administration and supervision: AM, WH and IM. Investigation, validation and visualisation: GM, ST and ES. Formal analysis and Data curation: GM, WH and VU. LM and AL made libraries with assistance from JL. Writing original draft: AM and GM. Writing review and editing: all authors.

## Competing Interests statement

The authors declare no competing interests.

## Notes

### Competing Interest Statement

The authors have declared no competing interest.

